# Transfer learning to leverage larger datasets for improved prediction of protein stability changes

**DOI:** 10.1101/2023.07.27.550881

**Authors:** Henry Dieckhaus, Michael Brocidiacono, Nicholas Randolph, Brian Kuhlman

**Affiliations:** Department of Biochemistry and Biophysics, University of North Carolina School of Medicine, Chapel Hill, North Carolina, USA; Division of Chemical Biology and Medicinal Chemistry, University of North Carolina Eshelman School of Pharmacy, Chapel Hill, North Carolina, USA; Department of Bioinformatics and Computational Biology, University of North Carolina School of Medicine, Chapel Hill, North Carolina, USA; Lineberger Comprehensive Cancer Center, University of North Carolina School of Medicine, Chapel Hill, North Carolina, USA

## Abstract

Amino acid mutations that lower a protein’s thermodynamic stability are implicated in numerous diseases, and engineered proteins with enhanced stability are important in research and medicine. Computational methods for predicting how mutations perturb protein stability are therefore of great interest. Despite recent advancements in protein design using deep learning, *in silico* prediction of stability changes has remained challenging, in part due to a lack of large, high-quality training datasets for model development. Here we introduce ThermoMPNN, a deep neural network trained to predict stability changes for protein point mutations given an initial structure. In doing so, we demonstrate the utility of a newly released mega-scale stability dataset for training a robust stability model. We also employ transfer learning to leverage a second, larger dataset by using learned features extracted from a deep neural network trained to predict a protein’s amino acid sequence given its three-dimensional structure. We show that our method achieves competitive performance on established benchmark datasets using a lightweight model architecture that allows for rapid, scalable predictions. Finally, we make ThermoMPNN readily available as a tool for stability prediction and design.

## Introduction

Proteins are a diverse, useful class of molecules that have found roles in a variety of clinical, industrial, and research settings^1–3^. Thermodynamic stability is a key property of any engineered protein, in part because naturally evolved proteins are typically only marginally stable under ambient conditions^4^. This is often sufficient for evolutionary purposes but is frequently a limiting factor in the utility of recombinant proteins for therapeutics, biocatalysis, and research applications. As a result, chemical biology investigations often seek to identify advantageous point mutations that may stabilize the candidate protein. While experimental stability optimization can be achieved using directed evolution^5^, computational stability models offer the potential for faster, cheaper, and more easily scalable optimization protocols^4^.

Many *in silico* methods have been developed to predict the effect of point mutations on protein stability given an initial structure. Historically, most methods have used empirical energy functions that model covalent and non-covalent interactions between atoms to evaluate mutations^6–8^. Other widely used protocols also incorporate sequence-derived information such as a position-specific substitution matrix^9^ or evolutionary consensus scoring^10^. Recent efforts have attempted to apply deep learning to this problem, each with their own limitations. Convolutional neural networks (CNNs) such as ThermoNet^11^ and RaSP^12^ require voxelization and computationally expensive convolutions, while methods using more efficient graph neural networks (GNNs) such as ProS-GNN^13^ and BayeStab^14^ require generation of a modelled mutant structure for inference, adding substantial runtime and introducing opportunity for error and model bias. There is therefore a need for fast, robust *in silico* methods to predict changes in thermodynamic stability (ΔΔG°) of potential point mutations given an initial wildtype structure.

Recent achievements using large language models (LLMs) for protein structure prediction have inspired models using pre-learned sequence embeddings to train models for various protein design tasks via transfer learning^15^, including for sequence-based stability prediction^16,17^. At the same time, Dauparas et al. released ProteinMPNN, a message-passing neural network (MPNN) trained on 19,700 protein clusters comprising the entire Protein Data Bank (PDB) (after quality filtering) to recover native-like sequences from a given protein backbone^18^. To achieve this goal, ProteinMPNN predicts the probability of all 20 amino acids being the native residue for a given position based on learned structural patterns found in natural proteins from the PDB. Native protein sequences with known structures are assumed to be at least marginally stable under normal conditions, meaning that the amino acid probabilities predicted by ProteinMPNN should correlate with the relative stabilities of the corresponding point mutants. However, evolution seldom optimizes for stability alone. Other properties such as solubility and enzymatic activity may necessitate compromises when evolving native sequences, and even when they do not, protein stability has diminishing returns beyond the bare minimum needed to ensure survival under ambient conditions^4^. As a result, using ProteinMPNN to naively predict ΔΔG° values is likely to be insufficient to achieve competitive performance. We therefore hypothesized that a model such as ProteinMPNN trained on sequence recovery may be amenable to transfer learning to enable accurate ΔΔG° prediction.

Our method, ThermoMPNN, takes advantage of knowledge overlap between the tasks of sequence recovery and stability optimization by using pre-trained ProteinMPNN embeddings as input features for transfer learning. In doing so, we effectively leverage two complementary large-scale datasets: the sequence recovery dataset used to train ProteinMPNN and a new “Megascale dataset” that includes experimental stability measurements on hundreds of thousands of mutations from ∼300 proteins^19^. Until recently, the most comprehensive datasets available for training and evaluating protein stability predictors were compilations from separate studies, and the number of cataloged mutations were frequently on the scale of less than 10,000 mutations. Due to the combination of transfer learning and training with the Megascale dataset, ThermoMPNN learns generalizable structural determinants of stability and achieves competitive performance on a variety of benchmarks. We also investigate different training regimes to determine the contribution of each dataset and model component. Finally, we profile the prediction patterns and biases of ThermoMPNN in comparison with its parent sequence recovery model.

## Results

### ThermoMPNN Architecture

ThermoMPNN (Fig. 1A) consists of two modules: a pre-trained ProteinMPNN model^18^ and a novel stability prediction module. As input features, ProteinMPNN uses pairwise distances between the backbone atoms of the target residue and the 48 nearest neighboring residues, encoded as Gaussian radial basis functions (RBFs). When used as a component of ThermoMPNN, the wild type (WT) amino acid sequence of the protein is also passed as a feature to ProteinMPNN via the decoding mechanism. ProteinMPNN (Fig. 1A, left box) is a graph neural network that includes three encoder layers (light blue box) followed by three decoder layers (light gold box). During encoding, message passing between nodes (one node for each residue) and edges (each residue pair is connected by an edge) allows each residue to learn about its structural environment. The decoder layers incorporate the WT sequence embedding along with node and edge embeddings to predict favorable amino acids for selected residues in the input structure, with intermediate information stored in 128-dimensional embeddings in each decoder layer (dark gold boxes). In ThermoMPNN, instead of passing predictions all the way through to the sequence recovery output layer, we extract these decoder embeddings from each layer, keeping only the embeddings for the residue that is being mutated (purple bars). This is then concatenated with the corresponding sequence embedding (green box) to produce the input vector for the stability prediction module (Fig. 1A, right box).

**Figure 1:**
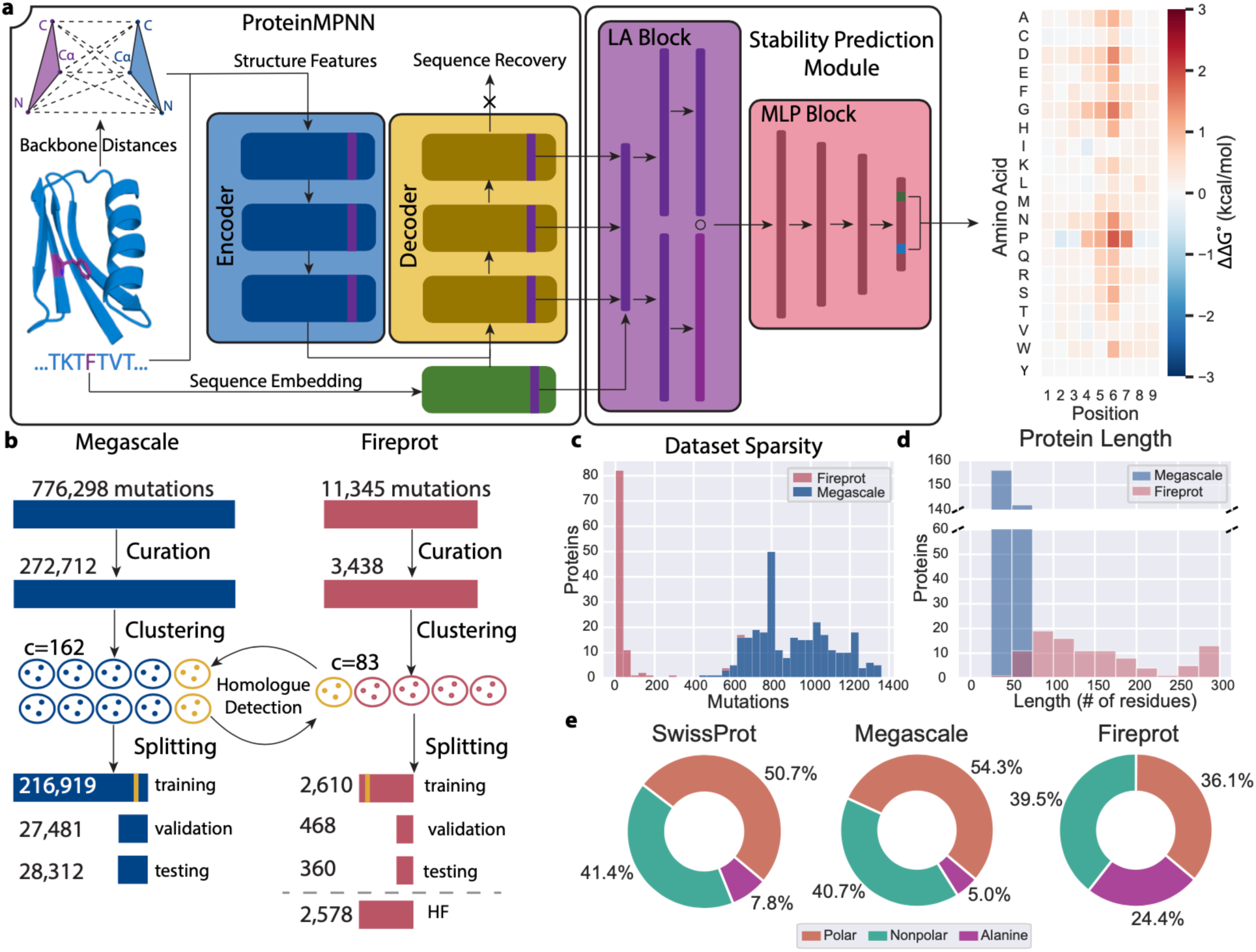
ThermoMPNN architecture and primary dataset statistics. **(a)** Model architecture of ThermoMPNN, a graph neural network trained on embeddings extracted from a pre-trained sequence recovery model (ProteinMPNN, left panel) to predict thermostability changes caused by protein point mutations. The input protein is passed through ProteinMPNN, where the learned embeddings from each decoder layer are extracted and concatenated with the learned sequence embedding to create a vector representation of the residue environment. This vector is passed through a light attention block (LA, purple block) which uses self-attention to reweight the vector based on learned context. Finally, a small multilayer perceptron (MLP, red block) predicts a ΔΔG° for mutation to each possible amino acid. **(b)** Curation, clustering, and data splitting procedure for the Megascale and Fireprot datasets used in this study. Each split is labelled with its total number of mutations, and homologues are shown in yellow. Each clustering result is labeled with the number of clusters in each dataset. **(c)** Histogram of mutations per protein distribution for each dataset. **(d)** Histogram of protein length distribution for each dataset. **(e)** Donut charts of percentage of mutations to alanine compared to other polar and nonpolar residues for each dataset, along with natural residue abundance for all proteins in the SwissProt database for comparison^24^.

The first component of the stability prediction module is a light attention (LA) block (Fig. 1A, purple block). This allows ThermoMPNN to reweight the input vector using contextual information via self-attention. Light attention has recently been shown to improve sequence-based protein localization^15^ and ΔΔG° prediction^16^ from LLM sequence embeddings, but this work is the first to our knowledge to utilize light attention for refinement of structural embeddings. The adjusted embedding is then passed through a small multi-layer perceptron (MLP) with two hidden layers (Fig. 1A, red block) to produce a final predicted ΔΔG° value.

### Dataset Preparation

The Megascale dataset used in this study was derived from a recent study by Tsuboyama et al. in which point mutation ΔΔG° values were inferred from protease sensitivity experiments^19,20^. This approach enables derivation of reasonably accurate ΔΔG° values at mega-scale (776,000 data points), which is multiple orders of magnitude larger than any previously utilized thermodynamic stability dataset derived from low-throughput biophysical assays (<10,000 data points). The ΔΔG° values must be inferred using a kinetic model, but they correlate well with previous ΔΔG° measurements, with observed Pearson correlations between 0.75-0.96^19^. For this study, data curation (Fig. 1B) was performed by selecting only single point mutations with reliable ΔΔG° values and valid wildtype structures to obtain a final dataset of 272,712 mutations across 298 proteins (Fig. 1B, blue). To evaluate model performance on thermodynamic stability data gathered with more traditional biophysical techniques, we aggregated an additional dataset by matching sequence-based stability data in the FireProtDB database^21^ to experimental structures in the PDB^22^. Data curation consisted of removing duplicates and points with missing information, then selecting the measurement for each mutation nearest to biological pH. The final dataset, which we will refer to as the Fireprot dataset, consisted of 3,438 mutations for 100 unique proteins (Fig. 1B, red). It should be noted that this dataset is a similar size to common literature training sets (e.g., S2648, Q3421, Q3488) with significant overlap in protein identity.

Both curated datasets were then clustered using MMseqs2^23^ with a stringent sequence identity cutoff of 25%. The two datasets were then cross-referenced to detect any homology overlap between the datasets (Fig. 1B, yellow), and any protein clusters with a homology match were automatically assigned to their respective training dataset. This ensured that none of the proteins in either Megascale or Fireprot test sets have any homologues in either training set. The remaining clusters were then randomly split to produce an approximately 80/10/10 split of mutations for each dataset. When splitting the Fireprot dataset, proteins with >250 data points were also automatically assigned to the training set, since their inclusion in the validation or test sets could have dominated any performance estimation, skewing the results. Finally, a fourth split, “homologue-free” (HF) Fireprot, was made for evaluating models trained solely on Megascale data, by retaining all Fireprot data except for those overlapping with Megascale data.

### Dataset Statistics

The Megascale and Fireprot datasets used in this study presented multiple key differences that contribute to their combined utility for assessing model performance (Fig. 1C-E). First, all Megascale proteins were represented by at least 400 measurements (Fig. 1C), with a maximum of around 1,400 measurements representing near-total site-saturation mutagenesis coverage of a miniprotein (20 amino acids × 70 residues = 1,400 measurements). On the other hand, 85 of 100 Fireprot proteins had less than 50 measurements each, while just a few well-studied proteins (streptococcal protein G and staphylococcal nuclease) comprised 1/3 of the dataset.

The Megascale dataset was also populated entirely by smaller proteins (<75 residues) (Fig. 1D) due to the restrictions imposed by oligonucleotide library synthesis. While the Fireprot dataset included some small proteins of a similar length (12 proteins of <75 residues), its members demonstrated both a greater average length and a wider distribution of lengths. The Fireprot dataset also exhibited a significant bias toward mutations to alanine (Fig. 1E), likely due to the popularity of alanine sweeps in mutagenesis experiments. This alanine bias was at the expense of mutations to polar residues, which comprise the majority (54.3%) of the Megascale dataset but only 36.1% of the Fireprot dataset. As a result, the mutation composition of the Megascale dataset better resembles the composition of naturally occurring proteins, as calculated from all proteins in the SwissProt database^24^.

### Ablation Study

To determine key contributions to ThermoMPNN’s performance, we conducted an ablation study by removing several components of the pipeline (Table 1). We found that using the published, pre-trained ProteinMPNN model alone for rank-ordering mutations was reasonably effective, with a Spearman correlation (SCC) of 0.487 ± 0.006 for the Megascale dataset and 0.50 ± 0.01 for the Fireprot dataset. The transfer learned ThermoMPNN model improved substantially over this baseline, with corresponding SCCs of 0.725 ± 0.003 and 0.657 ± 0.003, respectively. Importantly, starting from the pre-trained ProteinMPNN weights optimized for sequence recovery training was crucial for obtaining strong performance, with a significant drop-off observed (SCCs of 0.642 ± 0.005 and 0.50 ± 0.02) when training the model from naïve weights compared to transfer learning. However, attempting to optimize the pre-trained ProteinMPNN weights via fine-tuning produced mixed results. Although fine-tuning improved scores on the Megascale test set (SCC of 0.747 ± 0.001), this failed to translate to the Fireprot dataset, with mixed results compared to the transfer learning regime. Moreover, significant overfitting was also observed when fine-tuning (Supplementary Fig. 1). For these reasons, we opted to use transfer learning rather than fine-tuning for all further experiments. The LA module produced a small but consistent performance boost across both datasets, with no overfitting observed despite using a greater number of parameters (2.7M) than ProteinMPNN (1.7M).

**Table 1:** ThermoMPNN ablation study results (mean ± s.d.) on Megascale and Fireprot datasets.

| Dataset | Model | RMSE ↓ | PCC ↑ | SCC ↑ |
| --- | --- | --- | --- | --- |
| **Megascale (test)** | ProteinMPNN | $3.91 \pm 0.09$ | $0.43 \pm 0.01$ | $0.487 \pm 0.006$ |
| | ThermoMPNN | $0.708 \pm 0.007$ | $0.754 \pm 0.004$ | $0.725 \pm 0.003$ |
| | No light attention | $0.716 \pm 0.001$ | $0.742 \pm 0.002$ | $0.711 \pm 0.004$ |
| | No pre-training | $0.789 \pm 0.005$ | $0.689 \pm 0.005$ | $0.642 \pm 0.005$ |
|  | Added fine-tuning | **$0.69 \pm 0.01$** | **$0.780 \pm 0.002$** | **$0.747 \pm 0.001$** |
| **Fireprot (HF)** | ProteinMPNN | $3.4 \pm 0.1$ | $0.43 \pm 0.02$ | $0.50 \pm 0.01$ |
| | ThermoMPNN | **$1.51 \pm 0.01$** | **$0.650 \pm 0.005$** | $0.657 \pm 0.003$ |
| | No light attention | $1.562 \pm 0.006$ | $0.630 \pm 0.001$ | $0.634 \pm 0.001$ |
| | No pre-training | $1.66 \pm 0.04$ | $0.54 \pm 0.02$ | $0.50 \pm 0.02$ |
|  | Added fine-tuning | **$1.53 \pm 0.01$** | **$0.657 \pm 0.006$** | **$0.666 \pm 0.004$** |

### Effect of Training Data on ThermoMPNN

We next examined the effect of different training datasets on ThermoMPNN performance (Fig. 2A) in order to discriminate between contributions of model architecture and dataset composition. We found that ThermoMPNN metrics were significantly degraded when trained on the Fireprot dataset, with Pearson correlations (PCCs) of 0.49 ± 0.04 and 0.35 ± 0.02 on the Megascale and Fireprot test sets, respectively, compared to 0.754 ± 0.004 and 0.53 ± 0.005 when trained on the Megascale dataset. Two methods of co-training with both datasets were explored: concurrent training, wherein each epoch included batches from both datasets, and sequential training, wherein training on Megascale was followed by separate training epochs on Fireprot. Neither of these methods yielded any improvement over Megascale-only training. To determine what aspect(s) of Megascale data were most important for this training boost, the Megascale dataset was randomly subsampled to create a version with an equal number of proteins (Megascale SP) and a version with an equal number of total mutations (Megascale SM) to the Fireprot dataset. Models trained on each of these subsampled datasets also outperformed the Fireprot dataset when evaluated on the Megascale test set, although Megascale SM had similar performance on Fireprot test set. Notably, the Fireprot test set was more challenging to predict than the Megascale test set for all models tested.

**Figure 2:**
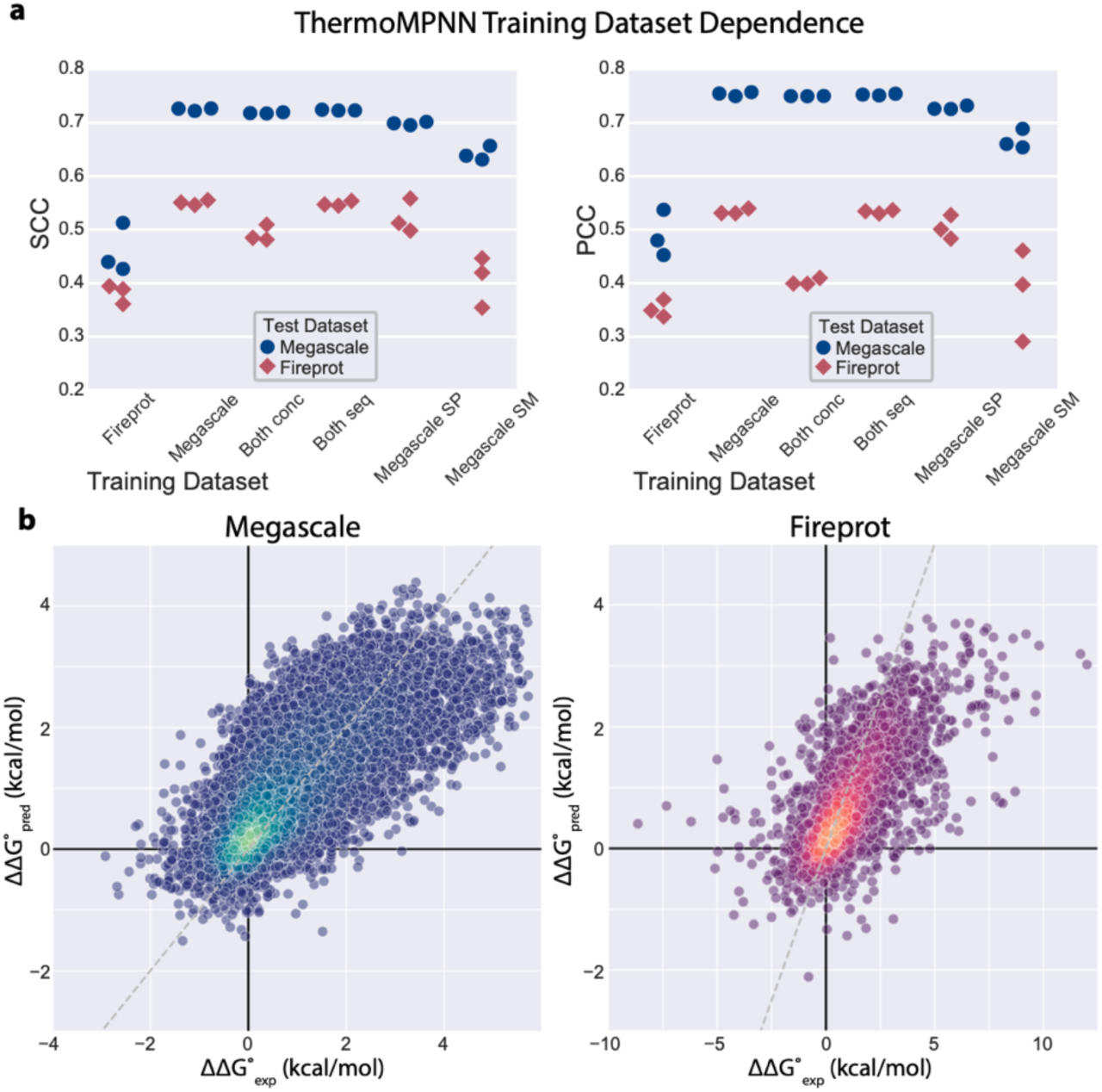
ThermoMPNN performance analysis. **(a)** Spearman (SCC) and Pearson correlation coefficients (PCC) calculated on predictions for models trained (in triplicate) using different datasets on Megascale (blue circle) and Fireprot (red diamond) test datasets. Models were also evaluated after either concurrent (Both conc) or sequential (Both seq) co-training, as well as training on only a subset of Megascale data including a similar number of proteins (Megascale SP) or a similar number of mutations (Megascale SM) to the Fireprot data. **(b)** ThermoMPNN predictions on the Megascale test (left, N = 28,312) and Fireprot HF (right, N = 2,578) datasets plotted vs their respective experimental values with identity line (dashed grey line) for reference. Points are colored by local density of mutations (lighter = denser) using a fitted Gaussian kernel density estimate (KDE) function.

Figure 2B shows the predictions of the top performing ThermoMPNN model on the Megascale (test) and Fireprot (HF) datasets plotted against their respective experimental measurements. One important note is that the Megascale data spans a substantially smaller dynamic range (-3 to 5 kcal/mol) than that of the Fireprot data (-9 to 12 kcal/mol). This means that ThermoMPNN, trained on the Megascale dataset, struggles to accurately predict ΔΔG° for outlier mutations. However, most Fireprot mutations (96.7%) fall within the dynamic range of the Megascale dataset, for which a fair correlation can be observed between predicted and experimental ΔΔG°. These findings indicate a potential limitation of the Megascale dataset as a tool for training stability predictors that are not explicitly built to extrapolate beyond the dynamic range of the training set.

### Comparison of ThermoMPNN with Other Models

We next conducted several comparisons to benchmark ThermoMPNN against other published models (Fig. 3 and Table 2). First, we tested different published methods with readily accessible code against our Megascale and Fireprot datasets (Fig. 3A): Rosetta (stabilize_pm protocol)^25^, 2-stage 3D CNN model RaSP^12^, and sequence-based transformer PROSTATA^17^. We found ThermoMPNN to be the most robust predictor that we evaluated on these two datasets, with a Pearson correlation 0.04-0.05 higher than any other method on each dataset. Re-training on the Megacsale dataset significantly improved both RaSP and PROSTATA scores on the Megascale test set but did not cause any improvement on the Fireprot HF set (Supplementary Table 2).

**Figure 3:**
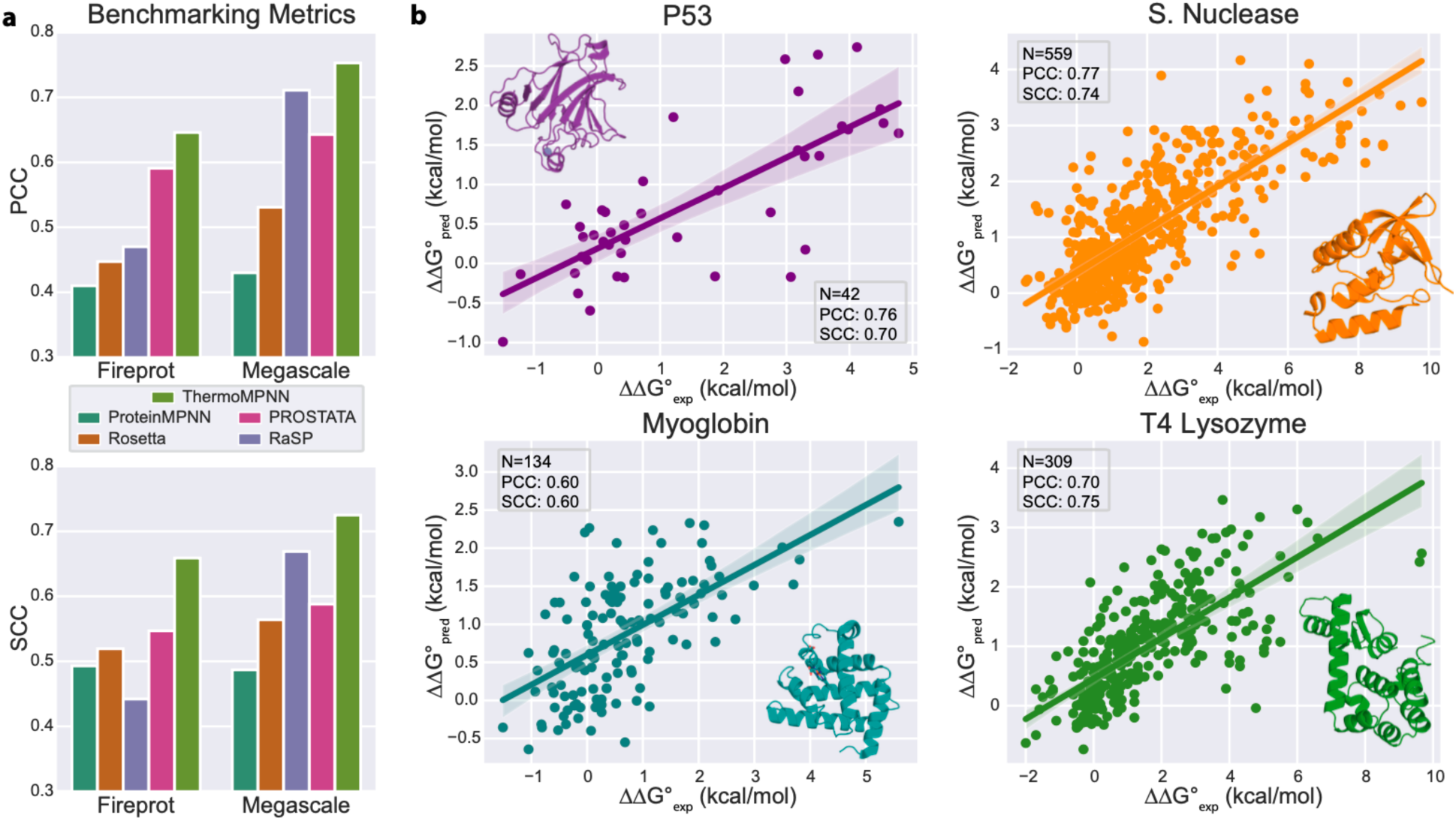
Literature comparison and case studies for ThermoMPNN predictions. **(a)** Comparison of Spearman (SCC, top) and Pearson (PCC, bottom) correlation coefficients for selected literature methods vs ThermoMPNN on Fireprot (HF) and Megascale (test) sets. **(b)** Scatter plots of stability change predictions (ΔΔG°_pred_) vs experimental measurements (ΔΔG°_exp_) for external test proteins p53 (PDB ID: 2OCJ), staphylococcus nuclease (1EY0), myoglobin (1BZ6), and T4 lysozyme (2LZM) with fitted regression line (shaded region = 95% confidence interval) and key statistics (N = number of mutations, PCC = Pearson correlation coefficient, SCC = Spearman correlation coefficient). The cartoon representation of each protein is included for reference.

**Table 2:**
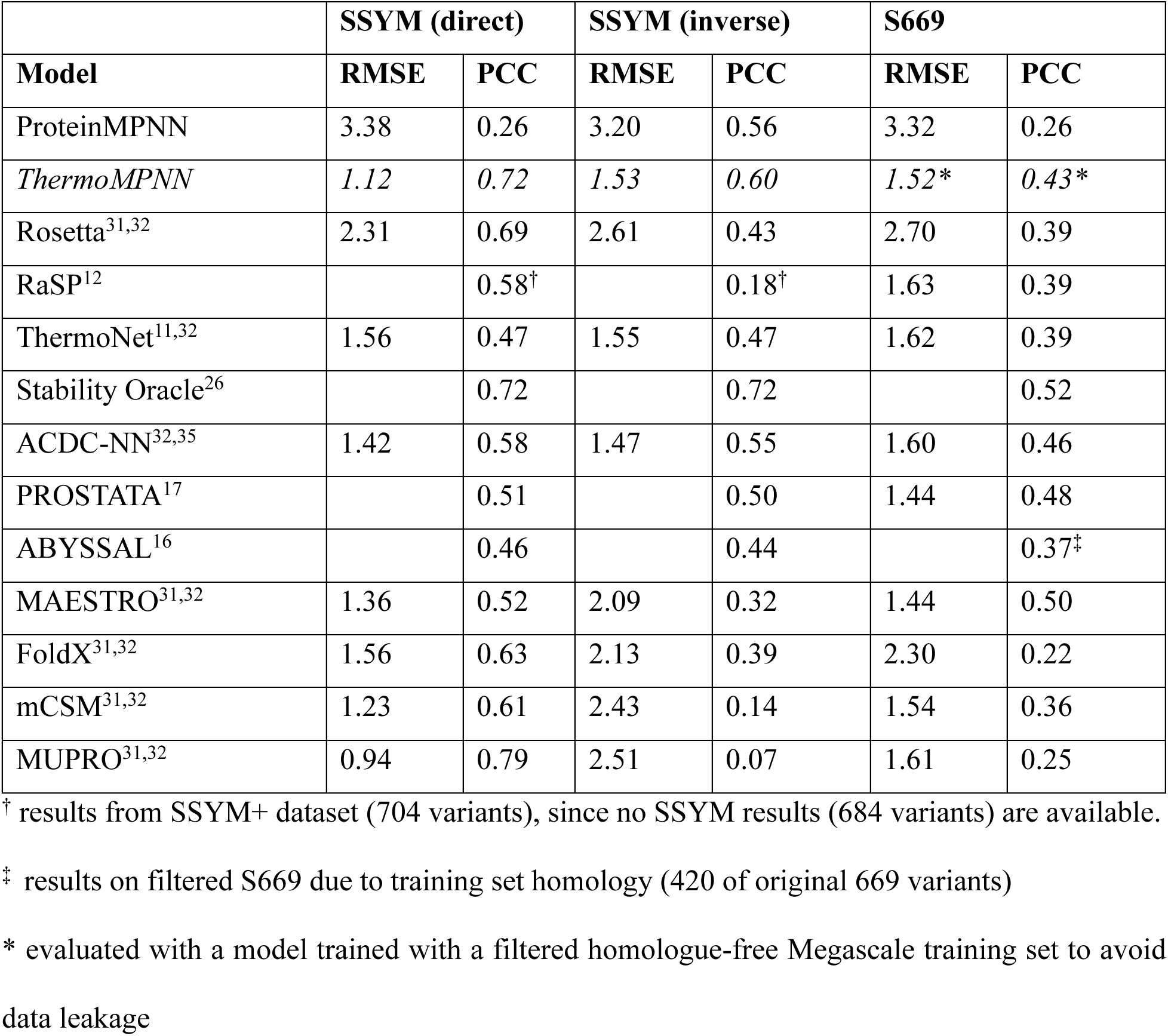
ThermoMPNN comparison with literature methods on SSYM and S669 datasets.

To further evaluate the performance of ThermoMPNN on unseen test data, we performed predictions for the SSYM and S669 datasets (Table 2). We found that ThermoMPNN achieves state-of-the art performance on both direct and inverse mutations from SSYM, albeit with higher absolute scores on the direct mutations (PCCs of 0.72 and 0.60, respectively). Only one prior method, recently released graph transformer Stability Oracle, outperformed ThermoMPNN on SSYM inverse mutations (PCC = 0.72), also achieving the same score (PCC = 0.72) on direct mutations. The code for this method is not yet publicly available for further comparison with ThermoMPNN on our own datasets. Consistent with previous works, the S669 dataset was more challenging for our model as well, with a PCC of 0.43 outperforming many structure-based methods (RaSP, Rosetta, etc.) but falling short of some other recent efforts (PROSTATA, ACDC-NN, etc.). Next, we evaluated ThermoMPNN in detail on four case studies (Fig. 3B), including common literature examples P53 and Myoglobin, as well as the two Fireprot-HF dataset proteins with the most mutations (staphylococcal nuclease, PDB ID: 1EY0; and T4 lysozyme, PDB ID: 2LZM). We found ThermoMPNN performed well on all these examples, with Myoglobin producing the lowest PCC of 0.60, compared to between 0.70 and 0.77 for the other three proteins.

### Exploration of ThermoMPNN Prediction Trends

We next performed 5-fold cross-validation on the Megascale dataset and analyzed the aggregated ThermoMPNN predictions obtained for all Megascale and Fireprot (HF) proteins (N = 387). Comparison of mutation preferences for ThermoMPNN against ProteinMPNN (Fig. 4A) revealed that ThermoMPNN significantly favors mutations toward hydrophobic residues, including on the protein surface. The largest individual changes were increased preferences for isoleucine, tryptophan, and surface arginine, along with reductions in surface lysine and glutamic acid residues. Examination of per-protein prediction root-mean-squared error (RMSE) (Fig. 4B) found that most proteins with <100 residues maintained an RMSE below 1.0 kcal/mol, while larger proteins were more likely to produce higher RMSEs. Despite this, 13 of 17 proteins with >100 residues still achieved an RMSE below 1.5 kcal/mol. De novo proteins achieved slightly lower average RMSE and a narrower score distribution than natural proteins, but there were also no large de novo proteins in our datasets, biasing a direct comparison. By comparison, natural proteins (up to 63 residues, max de novo protein length) had only slightly elevated RMSE (0.66) compared to the de novo proteins (0.60).

**Figure 4:**
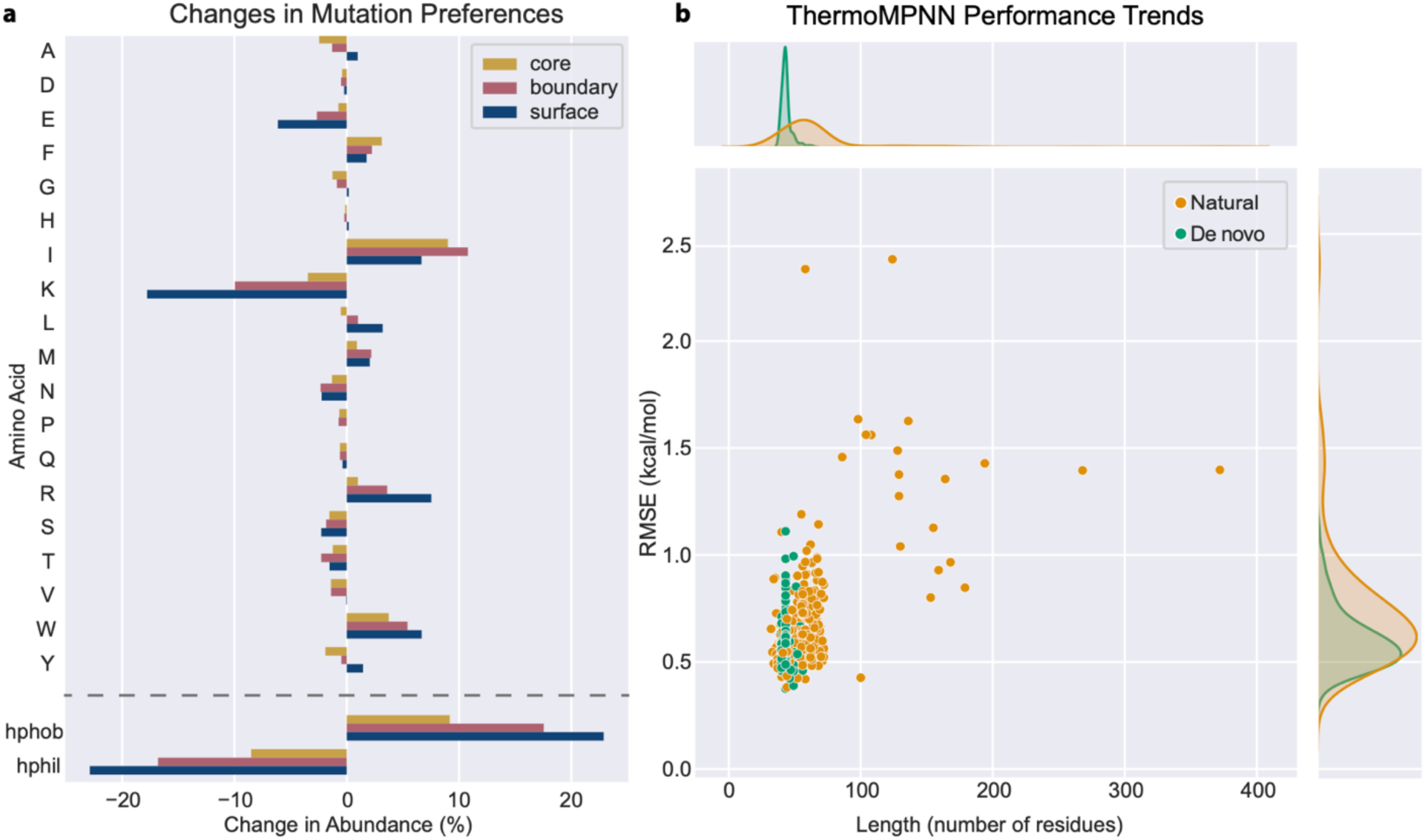
Exploration of ThermoMPNN prediction preferences and performance trends. **(a)** Change in amino acid preferences of ThermoMPNN compared to ProteinMPNN if the most favorable mutation is chosen at every position for all Megascale and Fireprot (HF) proteins. Cysteine mutations were excluded from this analysis due to potential disulfide formation biases (Supplementary Fig. 2). **(b)** ThermoMPNN root-mean-squared error (RMSE) for all Megascale and Fireprot (HF) proteins (min. 25 mutations, N = 320 proteins) plotted by protein length, with joint kernel density estimate (KDE) plots of the length (top) and RMSE (right) distributions of natural (N = 211) and de novo (N = 109) proteins.

## Discussion

Our results support our initial hypothesis that models trained for sequence recovery can be utilized via transfer learning to achieve accurate prediction of thermodynamic stability changes. We have demonstrated how ProteinMPNN, itself a weak ΔΔG° predictor, can be combined with a relatively simple stability prediction module to obtain a robust ΔΔG° model using only structural features. Our optimized model, ThermoMPNN, achieved competitive performance both on our own curated datasets and on established literature benchmarks, with particularly strong performance on the symmetrical SSYM dataset (PCC = 0.72/0.60 on direct/inverse mutations) and common case study P53 (PCC = 0.76).

One advantage of a transfer learning approach is that we can leverage a far larger dataset than those available for thermodynamic stability modeling. The PDB-derived dataset used to train ProteinMPNN includes 19,700 protein clusters^18^, which is 100 times larger than even the new Megascale stability dataset (162 clusters). This enables training of larger, deeper models such as ProteinMPNN that would be overfit on any stability dataset. Indeed, we found that naïvely re-training ThermoMPNN for ΔΔG° prediction led to severe overfitting, even on the Megascale dataset. This may also explain why fine-tuning the entire ProteinMPNN model from pre-trained weights failed to improve ThermoMPNN predictions. By transfer learning with only the lightweight ThermoMPNN stability prediction module, we were able to avoid overfitting and fully leverage the available data to learn generalizable structural patterns. At the same time, the introduction of the new Megascale dataset represents a significant opportunity for training deep neural networks for ΔΔG° prediction, as demonstrated in the training dataset exploration section. As expected, Megascale-trained models outperformed those trained on the sparse, unbalanced Fireprot dataset, and that advantage was more dependent on the total number of mutations included (sparsity) than on the number of unique proteins in the training dataset. Another recent structure-based transfer learning method, Stability Oracle, observed similar performance boosts from both pre-training for sequence recovery and transfer learning using the larger, more robust Megascale dataset^26^.

A few potential limitations of the Megascale dataset as a machine learning training dataset became apparent during our analysis. First, the dynamic range of the proteolysis assay is limited to ∼5 kcal/mol^19^, while experimental stability datasets such as our Fireprot dataset may include mutations with up to ±10 kcal/mol ΔΔG°. This means models trained on Megascale have limited capability to predict large changes in stability, a property that we also observe in other recently published models utilizing the Megascale dataset^16,26^. Second, we found that surface mutations to cysteine were often observed to be highly stabilizing in the Megascale dataset, such that ThermoMPNN would heavily favor surface cysteine mutations unless omitted from the permitted residue options (Supplementary Fig. 2). This phenomenon is a known artifact of the assay in which inter-molecular disulfide bonds are introduced, disrupting detection of backbone cleavage events^27^. More generally, the theoretical maximum performance of models trained on proteolysis-derived ΔΔG° values remains indeterminate, since individual protein correlations with biophysical measurements range from near-perfect (0.96) to merely good (0.75)^19^.

We also examined how ThermoMPNN amino acid preferences differ from those of its parent model, ProteinMPNN. Models trained to prioritize thermodynamic stability typically bias toward hydrophobic mutations, particularly on the protein surface^28^, while sequence recovery models must balance stability with solubility, function, and other constraints. ThermoMPNN predictions follow this trend, shifting significantly toward hydrophobic surface mutations when compared to ProteinMPNN. The increased preference of ThermoMPNN for surface hydrophobic residues is consistent with directed mutagenesis studies that show that surface hydrophobics can bury appreciable nonpolar surface area to lower the free energy of folding^29^. Importantly, our model is similarly effective on both (small) natural and de novo designed proteins, since it does not rely upon evolutionary information (i.e., multiple sequence alignment). ThermoMPNN is also fast, capable of running site-saturation mutagenesis in just 2 seconds for a small (105 residue) protein and 8 seconds for a large (693 residue) protein on a single GPU (Supplementary Table 3). This could enable ThermoMPNN to be integrated into iterative *in silico* design protocols in the near future. To this end, we have released ThermoMPNN as both a standalone Python package and a web-accessible Colab notebook so that it may be useful to both developers and biologists (https://github.com/Kuhlman-Lab/ThermoMPNN). We hope that this work will serve as an example of the power of transfer learning for protein design objectives and the importance of leveraging all available data to enable widely applicable, readily repurposed models.

## Methods

### Dataset Curation

The Megascale dataset used in this study was derived from the updated April 2023 version released by Tsuboyama et al. (https://zenodo.org/record/7992926)^20^. The raw data was curated by removing points with ΔΔG° values marked unreliable, as well as all insertions, deletions, and multiple mutants. Proteins were then filtered to remove any data derived from perturbed wildtype measurements to avoid using inaccurate structures. This yielded a final Megascale dataset of 272,712 mutations across 298 proteins.

The Fireprot dataset used in this study was obtained by downloading the full FireProtDB database (https://loschmidt.chemi.muni.cz/fireprotdb/)^21^. Duplicate entries and those missing key data fields (ΔΔG°, PDB ID, UniProt ID, position, wildtype residue, and mutant residue) were removed, then the wildtype sequences were aligned to experimental structures from the PDB (https://www.rcsb.org/)^22^. Entries with multiple PDB IDs were manually disambiguated, oligomers were removed, and a small number of additional proteins were removed due to quality control issues (Supplementary Table 1). For mutations with multiple measurements, the entry obtained under pH closest to biological pH (7.4) was selected. The final Fireprot dataset consisted of 3,438 mutations across 100 proteins.

### Dataset Splitting

The MMseqs2 *easy-cluster* tool^23^ was used to cluster each dataset with a minimum sequence identity cutoff of 25%. This produced 163 clusters of between 1 and 27 members for the Megascale dataset and 83 clusters of between 1 and 3 members for the Fireprot dataset. The Mmseqs2 *easy-search* tool was used to detect homology matches between Megascale and Fireprot, again with a 25% identity cutoff. This returned 41 total matches between 32 Megascale and 11 Fireprot proteins. Random splitting produced 80/10/10 training/validation/testing splits containing 239/31/28 proteins for Megascale and 57/15/28 proteins for Fireprot (the homologue-free split contained 89 proteins). The Fireprot proteins with >250 data points were streptococcal protein G (PDB ID: 1PGA), staphylococcal nuclease (1EY0), and T4 lysozyme (2LZM).

### Dataset Analysis

Amino acid abundances for natural proteins in the SwissProt database were obtained from the literature^24^ and totaled according to the following groups: polar (CDEHKNQRSTY), nonpolar (FGILMPVW), and alanine (A).

### ThermoMPNN Architecture

ThermoMPNN (Fig. 1A) is a graph neural network that treats the input protein as a connected graph of nodes (residues) and edges (distances). Its first module consists of a pre-trained ProteinMPNN network, which we treat as a feature extractor by freezing all parameters. Pairwise atomic distances are calculated between all backbone atoms (N, Cα, C, O, and Cβ), and the 48 nearest neighbors are collected for each residue. These distance features are passed through 3 message-passing encoder layers in which information is “passed” between connected nodes (i.e., nearby residues). The encoder is followed by a decoder, also comprised of 3 layers, which utilize features from the encoder as well as any available sequence embeddings to “decode” residues one-by-one for sequence recovery. In ThermoMPNN, the intermediate embeddings for the target mutation position stored in each decoder layer are extracted and concatenated with the sequence embedding for the wildtype residue. This produces a vector of size 128+(128×N), where N is the number of decoder layers included (we use N = 2 for all described experiments).

The next component of ThermoMPNN is a light attention block. First, two independent padded convolutions (size = 9, stride = 1) are performed on the 1-dimensional input vector. The feature convolution is fed through a dropout layer (probability = 0.25), while the attention convolution is rescaled with a softmax layer. These two outputs are then multiplied elementwise to produce a reweighted feature vector of the same size according to learned attention patterns. Finally, this vector is passed through an MLP with two hidden layers (sizes 64 and 32) to predict a ΔΔG° which is normalized by subtracting the predicted mutant ΔΔG° from a predicted wildtype ΔΔG° at the same position.

### ThermoMPNN Training

Unless otherwise stated, all ThermoMPNN models were trained using an AdamW optimizer (learning rate = 0.001, weight decay = 0.01) with mean squared error (MSE) loss for 100 epochs. The ProteinMPNN model “v48_020.pt” (48 nearest neighbors, 0.2Å training noise) was used for all knowledge transfer experiments. Each training batch consisted of all mutations for a given protein, and all batches were sampled in each epoch. After training, the best model was selected based on the highest Spearman correlation achieved on the validation set. Training ThermoMPNN on the Megascale dataset took approximately 18 hours on a single V100 GPU.

### Ablation Study

ProteinMPNN rank-order mutation scores were obtained by extracting the normalized log-probabilities for each mutation and multiplying them by -1. To fine-tune ProteinMPNN, the parameters were unfrozen and co-trained according to the procedure described above. A hyperparameter sweep (from 1 x 10^-2^ to 1 x 10^-6^) identified an optimal learning rate of 1 x 10^-4^ for the ProteinMPNN layers. All models were trained in triplicate using three different random seeds in order to calculate mean and standard deviation of the performance metrics.

### Training Dataset Experiments

ThermoMPNN concurrent co-training was implemented by training on all batches of Megascale and Fireprot training data on each epoch with all other hyperparameters held constant. For sequential co-training, 50 epochs of Megascale training were completed as per the above procedure. Next, the learning rate was decreased from 0.001 to 0.0001 and the light attention block was frozen to only permit the final MLP to further train. Then 50 epochs of training on Fireprot dataset were completed.

### Comparison with Literature Methods

Several literature methods were evaluated on our Megascale and Fireprot datasets. For our Rosetta benchmark, we used the recently detailed point mutation stabilization protocol^25^. Code for RaSP (https://github.com/KULL-Centre/_2022_ML-ddG-Blaabjerg) and PROSTATA were downloaded from their respective GitHub repositories (https://github.com/mitiau/PROSTATA). RaSP and PROSTATA were initially evaluated using the default trained weights released by their respective authors but were re-trained using our Megascale training dataset for direct comparison with ThermoMPNN (Supplementary Table 2). For PROSTATA, the best scoring model (or ensemble) was included in the analysis, and a symmetrized Megascale training dataset was used to allow training on maximum available data. For RaSP, only the 2^nd^ stage (“Downstream”) model was re-trained on Megascale data – the author-provided weights for the self-supervised representation model were used for all experiments. Performance metrics for literature methods on SSYM and S669 datasets were obtained from prior literature (citations included in Table 2).

### ThermoMPNN Mutation Preferences

Protein structure regions (core/boundary/surface) were calculated by labeling each residue according to how many nearby (within 10Å) neighbors were found (according to Cα-Cα distances). Residues with ≥20 neighbors were classified as core, those with ≤15 neighbors as surface, and those in between 20 and 15 neighbors as boundary.

### ThermoMPNN Runtime Analysis

ThermoMPNN runtime analysis (Supplementary Table 3) was performed by averaging 5 replicate runs for each protein on a single V100 GPU with 16 GB VRAM and 8 CPUs.

## Data Availability

The curated Fireprot dataset utilized in this study can be downloaded from the provided Zenodo data repository (https://doi.org/10.5281/zenodo.8169289)^30^. Scripts for preprocessing the Fireprot dataset starting from the raw FireProtDB data are also included in the ThermoMPNN GitHub repository. All other datasets can be obtained from the ProtDDG-Bench repository (https://github.com/protddg-bench/protddg-bench) or from their respective literature sources. This included SSYM^31^, S669^32^, P53^33^, Myoglobin^34^, and Megascale^20^ datasets.

## Code Availability

The code for ThermoMPNN is available at https://github.com/Kuhlman-Lab/ThermoMPNN. We also provide a Colab implementation of ThermoMPNN in the same repository (Supplementary Fig. 3).

## Acknowledgements

This work was supported by the NIH grant R35GM131923 (B.K.), and by the National Science foundation fellowship DGE-2040435 (N.R.). The authors would like to thank David F. Thieker for his assistance with the Rosetta stabilize_pm protocol.

## Author contributions

**H.D.:** Conceptualization, Methodology, Software, Data Curation, Visualization, Writing – Original Draft, Writing – Review & Editing. **M.B.:** Conceptualization, Methodology, Software, Writing – Review & Editing. **N.R.:** Conceptualization, Methodology, Software, Writing – Review & Editing, Supervision. **B.K.:** Conceptualization, Writing – Original Draft, Writing – Review & Editing, Supervision, Project Administration, Funding Acquisition.

