## Supplementary Information for "Transfer learning to leverage larger datasets for improved prediction of protein stability changes"

Henry Dieckhaus^1,2^, Michael Brocidiacono^2^, Nicholas Randolph^1,3^, Brian Kuhlman^1,3,4^*

^1^ Department of Biochemistry and Biophysics, University of North Carolina School of Medicine, Chapel Hill, North Carolina, USA

^2^ Division of Chemical Biology and Medicinal Chemistry, University of North Carolina Eshelman School of Pharmacy, Chapel Hill, North Carolina, USA

^3^ Department of Bioinformatics and Computational Biology, University of North Carolina School of Medicine, Chapel Hill, North Carolina, USA

^4^ Lineberger Comprehensive Cancer Center, University of North Carolina School of Medicine, Chapel Hill, North Carolina, USA

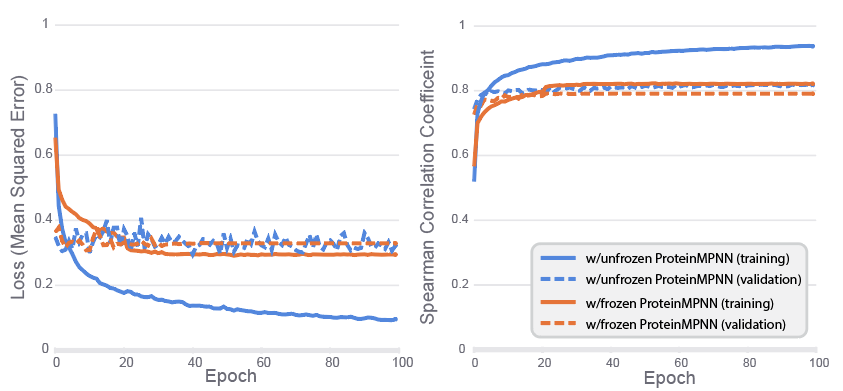

**Supplementary Figure 1: Training curves showing overfitting of ThermoMPNN model with unfrozen ProteinMPNN layers.** Training loss (left) and Spearman correlation (right) training (solid lines) and validation (dashed lines) curves for ThermoMPNN models trained on the Megascale dataset with frozen (orange) and unfrozen (blue) ProteinMPNN layers (learning rate of 0.0001 for unfrozen ProteinMPNN).

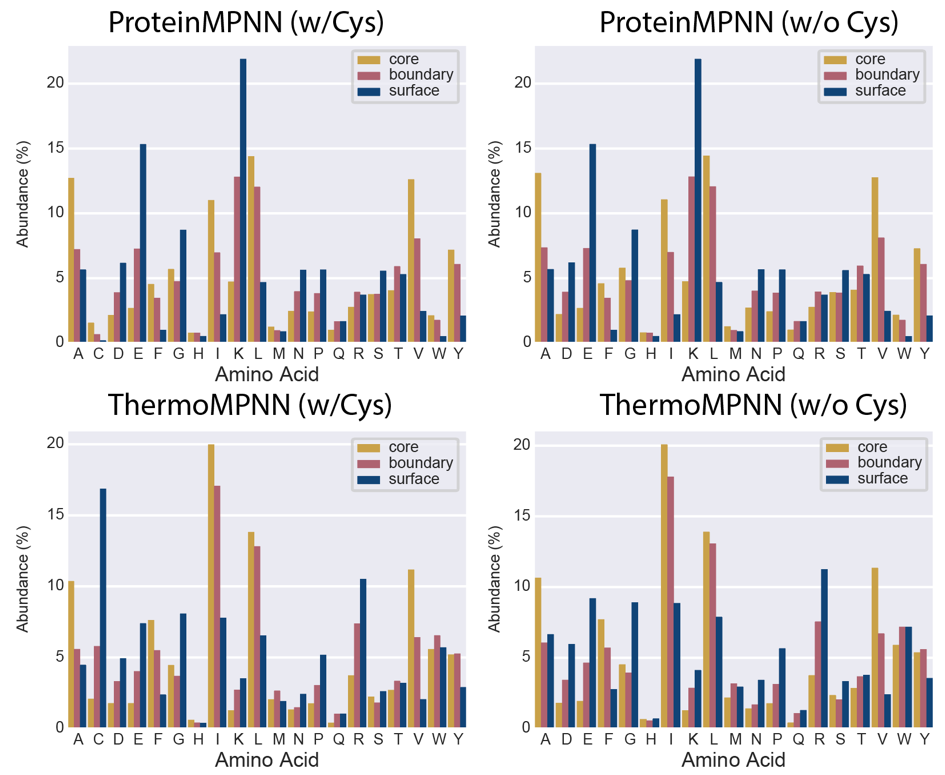

**Supplementary Figure 2: Amino acid preferences of ThermoMPNN and ProteinMPNN.** Plots of amino acid abundance of combined Megascale and Fireprot-HF datasets with (left) and without (right) allowing cysteine mutations.

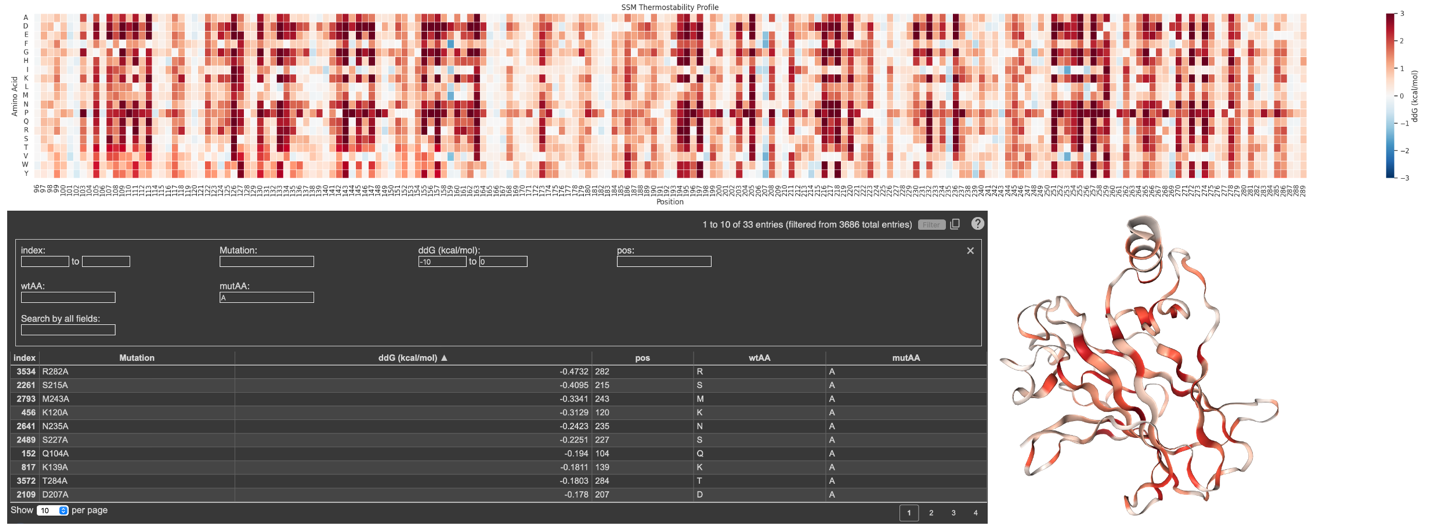

**Supplementary Figure 3: Example *in silico* mutagenesis results generated via the ThermoMPNN Colab notebook.** At top, a heatmap of all possible point mutations displays favored (blue) and disfavored (red) mutations for streptococcal protein G (PDB ID: 1PGA). At bottom left, an embedded table displays raw ΔΔG° values for all mutations and offers filtering and download capabilities. At bottom right, an interactive sensitivity map is generated by calculating the mean of the magnitude of all ΔΔG° values at a given position (darker red indicates greater mutation sensitivity).

**Supplementary Table 1: Manually Removed Fireprot Proteins**

| Protein Name | PDB ID | Reason for Removal | Data Points |
| --- | --- | --- | --- |
| SAC7D | 1C8C | Nucleotide bound | 3 |
| SSO7D | 1BF4 | Nucleotide bound | 25 |
| Ferric enterobactin | 1FEP | Disulfide formation | 7 |

**Supplementary Table 2: RaSP and PROSTATA Performance Before and After Retraining**

| Model | Training Data | Test Data | RMSE | PCC | SCC |
| --- | --- | --- | --- | --- | --- |
| PROSTATA | Megascale (training) | Megascale (test) | 0.834 | 0.643 | 0.588 |
| PROSTATA | Megascale (training) | Fireprot (HF) | 1.675 | 0.591 | 0.547 |
| PROSTATA | PROSTATA | Megascale (test) | 0.944 | 0.559 | 0.506 |
| PROSTATA | PROSTATA | Fireprot (HF) | 0.919^†^ | 0.874^†^ | 0.911^†^ |
| RaSP | Megascale (training) | Megascale (test) | 1.084 | 0.711 | 0.669 |
| RaSP | Megascale (training) | Fireprot (HF) | 1.861 | 0.470 | 0.442 |
| RaSP | RaSP | Megascale (test) | 1.029 | 0.631 | 0.568 |
| RaSP | RaSP | Fireprot (HF) | 1.713 | 0.480 | 0.467 |

^†^ Duplicate proteins and homologues were observed between PROSTATA training dataset and Fireprot dataset, likely inflating performance.

**Supplementary Table 3: ThermoMPNN Runtime Analysis on Example Proteins.**

| Protein | PDB ID (chain) | Residues | Wall-clock time (s)^†^ | | |
| --- | --- | --- | --- | --- | --- |
|  |  |  | Pre-processing | Inference | Total runtime  (per residue) |
| ELOB | 4AJY (B) | 105 | 0.01 | 2 | 0.02 |
| GCK | 4CDH (A) | 434 | 0.04 | 5 | 0.01 |
| F8 | 2R7E (A) | 693 | 0.07 | 8 | 0.01 |

^†^ Times are averaged across 5 replicate runs for each protein.
